# Individual differences in late positive potential amplitude and theta power predict cue-induced eating

**DOI:** 10.1101/2022.03.28.485549

**Authors:** Kyla D. Gibney, George Kypriotakis, Francesco Versace

## Abstract

Both the brain’s affective and cognitive systems are known to regulate cue-induced behaviors. Although it is established that individual differences in affective processing of food-related cues predict cue-induced eating, we have yet to investigate how both affective and cognitive mechanisms act in tandem to regulate cue-induced eating. We recorded electroencephalogram (EEG) from 59 adults while they viewed emotional & food-related images that preceded the delivery of food rewards (candies) or nonfood objects (beads). We measured the amplitude of the late positive potential (LPP) in response to images and power in the theta (4-8 Hz) frequency band after the candy or bead was dispensed to the participant. We found that individuals with larger LPP responses to food cues than to pleasant images (C>P group) ate significantly more during the experiment than did those with the opposite response pattern (P>C group, *p* < 0.001). Furthermore, we found that individuals with higher theta power after dispensation of the candy than of the bead (θCA>θBE) ate significantly more than did those with the opposite response pattern (θBE>θCA, *p* < 0.001). Finally, we found that the crossed P>C and θBE>θCA group ate less (*p* < 0.001) than did the other three groups formed by crossing the LPP and theta group assignments, who exhibited similar eating behavior on average (*p* = 0.662). These findings demonstrate that motivational salience and cognitive control converge to independently confer vulnerability or resilience to cue-induced behaviors, underscoring the need for individualized treatments to mitigate maladaptive behaviors.

**HIGHLIGHTS:**

- LPP responses to motivationally salient stimuli predict cue-induced eating.
- Theta power in the presence of food rewards predicts cue-induced eating.
- The LPP & theta may inform the development of personalized weight-loss interventions.

## 1. Introduction

Overweight and obesity, characterized by a body mass index of at least 25 kg/m^2^ and at least 30 kg/m^2^, respectively, increase the risk of cardiovascular disease, diabetes, and several types of cancer [1]. Losing even a modest amount of weight can have substantial health benefits, but most weight-loss interventions yield short-lived, suboptimal results [2,3]. Identifying the neurobiological mechanisms underlying excessive eating, the ultimate cause of weight gain [4,5], can help clinicians target the root causes of overeating, personalize interventions for weight loss, and improve weight loss treatment outcomes.

Neurobiological models of obesity have demonstrated that the brain’s reward and cognitive control systems both play a major role in regulating food intake [6,7]. The reward system guides eating behavior with “bottom-up” signals that dynamically assign motivational salience to food rewards and the cues associated with them [8,9]. In contrast, cognitive control systems exert “top-down” control over eating behavior by enabling the implementation of intentional, goal-directed behavior [10]. Failure of either mechanism can lead to maladaptive eating patterns, overeating of hyper-palatable foods, and weight gain [11].

Preclinical findings demonstrated that animals differ in their tendency to engage bottom-up versus top-down driven behaviors in the presence of cues signaling the impending delivery of food rewards (i.e., food-related cues) [12]. Individuals who attribute high motivational salience to food-related cues are prone to cue-induced compulsive reward-seeking behaviors [13]. On the other hand, those who do not attribute high levels of motivational salience to reward-related cues are less prone to cue-induced compulsive behaviors and are likely to implement goal-directed behaviors when faced with these cues [14].

We have demonstrated that humans are also characterized by individual differences in the tendency to attribute motivational salience to food-related cues [15] and that these differences underlie vulnerability to cue-induced eating [16]. In our experiments, we recorded event-related potentials (ERPs)—a direct measure of brain activity [17]—during a cued food delivery task. In this task, participants viewed emotional, neutral, and food-related images while we recorded electroencephalogram (EEG) from the scalp. After the presentation of some food-related images, we dispensed chocolate candies to the participants, which they could eat or discard [18]. To estimate the motivational salience of these images, we measured the amplitude of the late positive potential (LPP) in response to each image. The LPP is an ERP component that is reliably modulated by motivational salience: highly salient images such as erotica and mutilations prompt larger LPP responses than do images with lower salience, such as romantic or sad images [19–21]. We found that individuals with larger LPP responses to food-related cues than to pleasant images (C>P group) ate significantly more during the experiment than did those with larger LPP responses to pleasant images than to food-related cues (P>C group) [16].

These findings support the hypothesis that attributing higher motivational salience to food-related cues than to pleasant non-food-related stimuli increases vulnerability to cue-induced eating, but they are silent about the role of individual differences in the engagement of cognitive control systems in regulating cue-induced eating. Because results from animal models suggest that individuals who attribute high levels of motivational salience to food-related cues might also have poor top-down control over cue-induced behaviors [12,14,22,23], the present study aimed to elucidate how both cognitive and affective mechanisms act in tandem to regulate cue-induced eating.

Activity in the theta frequency band has been proposed as a reliable correlate of the engagement of higher cognitive functions [24,25]. Theta power over midfrontal scalp sites increases when participants engage cognitive control mechanisms to inhibit prepotent responses [26,27] or perform difficult tasks [28]. In light of these findings, we used theta power in an exploratory fashion to approximate the engagement of cognitive control systems in food-related decision-making during a cued food delivery task.

The present study aimed at investigating the role that individual differences in both the attribution of motivational salience to food-related cues and the engagement of cognitive control systems have in regulating cue-induced eating during the cued food delivery task. We expected to replicate our previous findings: namely, that individual differences in affective processing of cues predict cue-induced eating. We also aimed to elucidate whether the engagement of cognitive control systems—as indexed by theta power—differs between P>C and C>P individuals, or if cognitive control mechanisms contribute to cue-induced eating behavior irrespective of C>P and P>C status. Results demonstrating that midfrontal theta power differs between the C>P and P>C groups would suggest that individuals attributing higher motivational salience to food-related cues might also have difficulty engaging cognitive control mechanisms when making food-related decisions, in a manner similar to what has been observed in animal models. Meanwhile, results demonstrating that individual differences in midfrontal theta power predict eating behavior regardless of C>P vs and P>C status would suggest that the engagement of cognitive control systems regulates cue-induced eating independently from the tendency to attribute motivational salience to food-related cues. By elucidating whether motivational salience and cognitive control mechanisms converge to regulate cue-induced eating or do so independently, we may inform clinical researchers of effective mechanistic targets for weight loss and other clinical interventions aimed at reducing maladaptive, reward-seeking behaviors.

## 2. Materials and methods

### 2.1. Participants

We recruited sixty research participants from the Houston, TX, metro area using flyers and magazine and newspaper advertisements. Participants were eligible if they were 18 to 65 years of age, were neither pregnant nor breastfeeding, and did not have a history of psychiatric disorders, seizures, head injuries with loss of consciousness, uncorrected visual impairments, eating disorders, or allergies, or any other illnesses that would prevent them from eating chocolate candy. Participants received monetary compensation for their time and travel totaling up to $60 each. One participant was excluded from the final analysis due to incomplete data. Tables 1 and 2 show the demographic information for the participant sample.

**Table 1.**
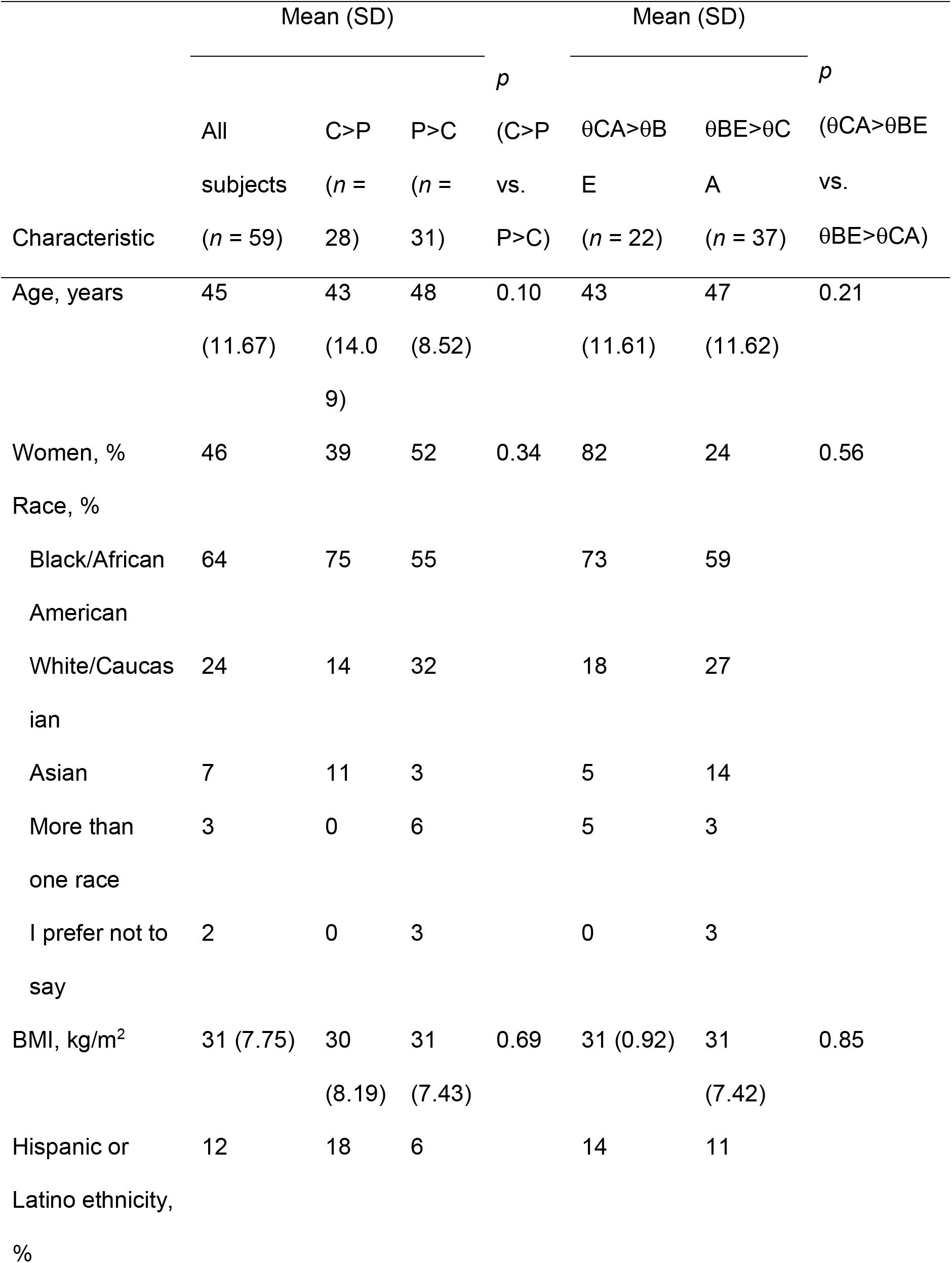
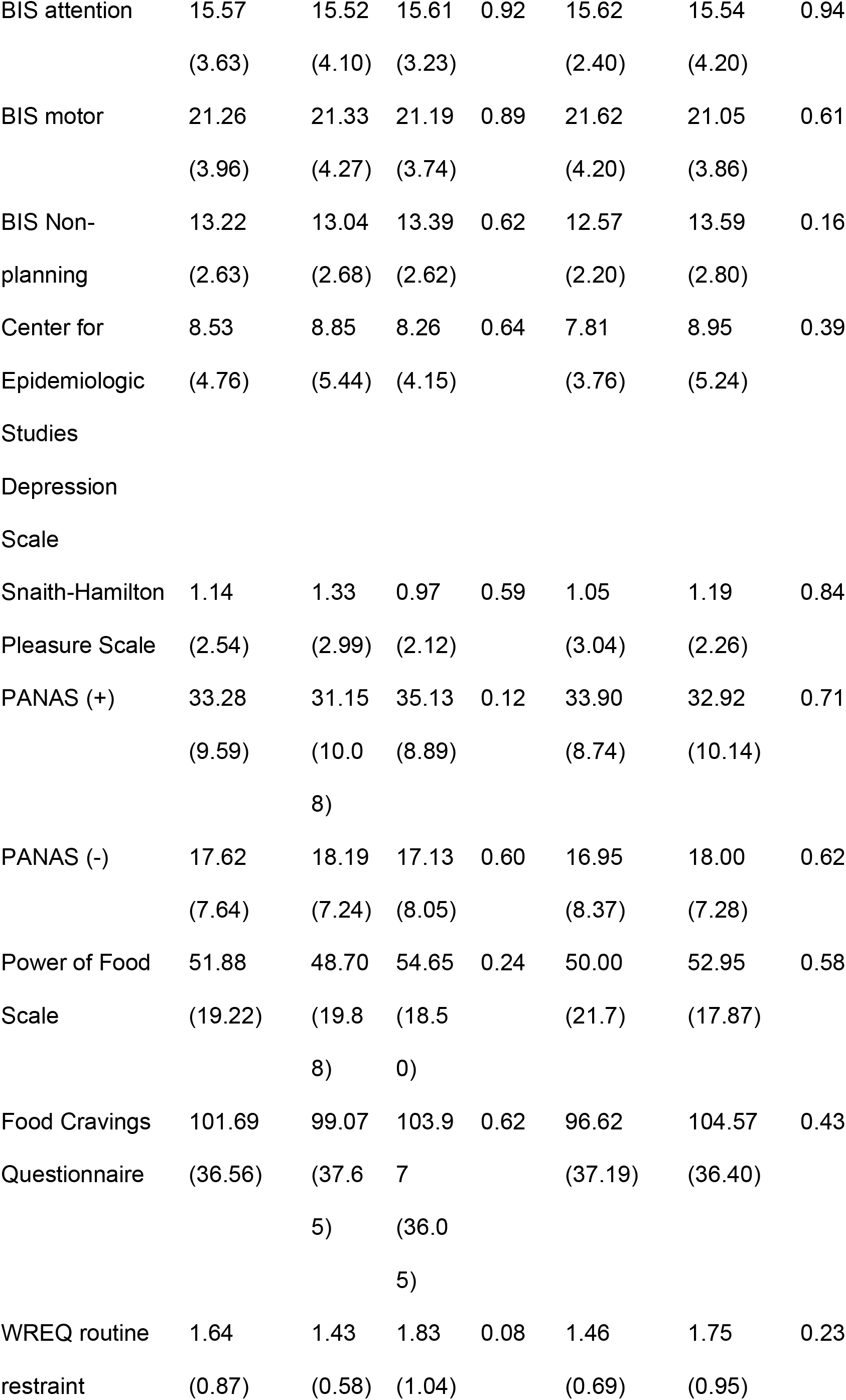
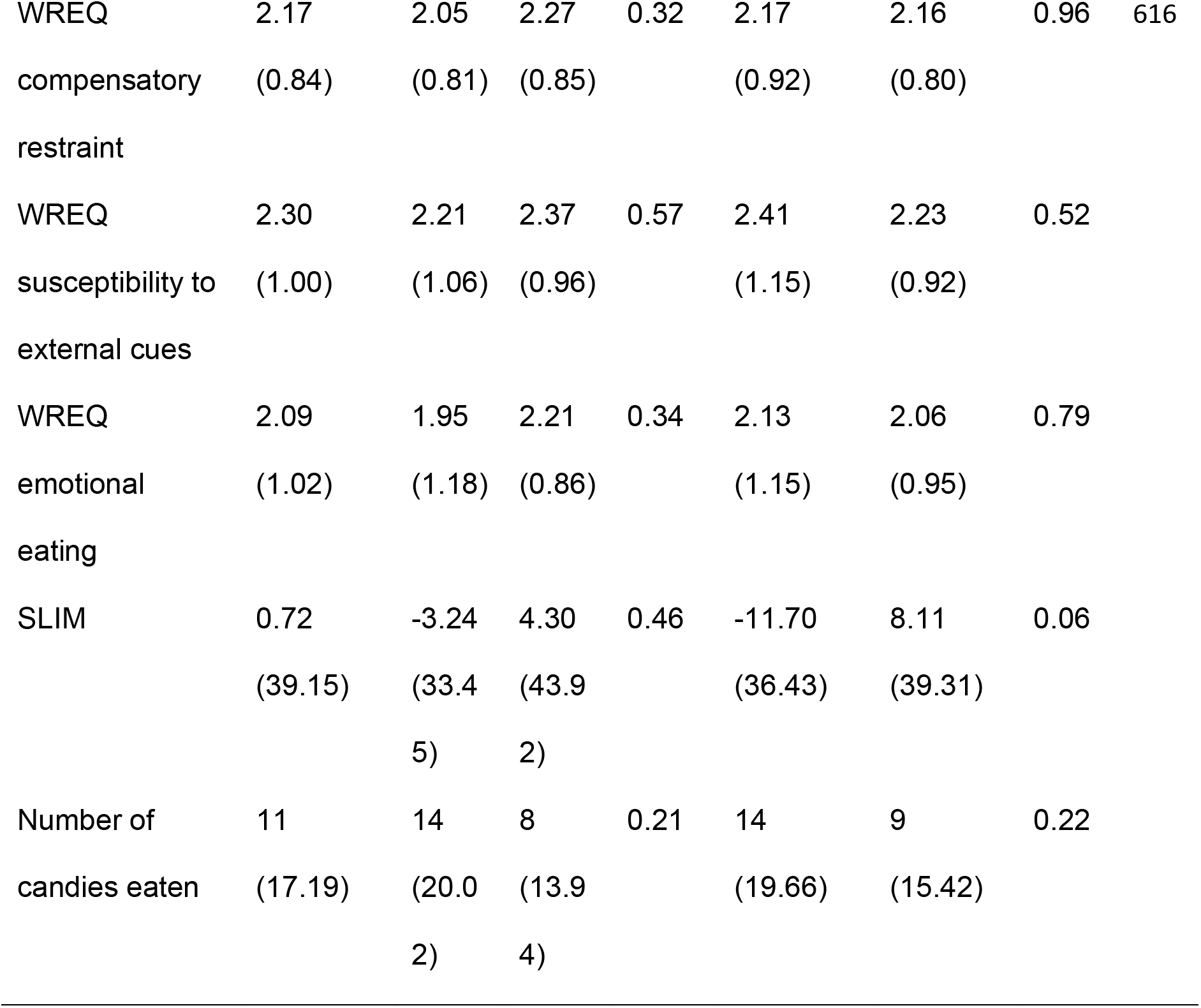
Demographic, biometric, and self-reported data for all subjects and the LPP and theta-based participant groups.

| Characteristic | Mean (SD) |  |  |  | Mean (SD) |  |  |
| --- | --- | --- | --- | --- | --- | --- | --- |
|  |  |  |  | *p* |  |  | *p* |
|  | All subjects<br>(*n* = 59) | C>P<br>(*n* = 28) | P>C<br>(*n* = 31) |  | θCA>θB<br>E<br>(*n* = 22) | θBE>θC<br>A<br>(*n* = 37) | (θCA>θBE<br>vs.<br>θBE>θCA) |
| Age, years | 45<br>(11.67) | 43<br>(14.09) | 48<br>(8.52) | 0.10 | 43<br>(11.61) | 47<br>(11.62) | 0.21 |
| Women, % | 46 | 39 | 52 | 0.34 | 82 | 24 | 0.56 |
| Race, % |  |  |  |  |  |  |  |
| Black/African American | 64 | 75 | 55 |  | 73 | 59 |  |
| White/Caucasian | 24 | 14 | 32 |  | 18 | 27 |  |
| Asian | 7 | 11 | 3 |  | 5 | 14 |  |
| More than one race | 3 | 0 | 6 |  | 5 | 3 |  |
| I prefer not to say | 2 | 0 | 3 |  | 0 | 3 |  |
| BMI, kg/m<sup>2</sup> | 31 (7.75) | 30<br>(8.19) | 31<br>(7.43) | 0.69 | 31 (0.92) | 31<br>(7.42) | 0.85 |
| Hispanic or Latino ethnicity, % | 12 | 18 | 6 |  | 14 | 11 |  |
| BIS attention | 15.57 | 15.52 | 15.61 | 0.92 | 15.62 | 15.54 | 0.94 |
|  | (3.63) | (4.10) | (3.23) |  | (2.40) | (4.20) |  |
| BIS motor | 21.26 | 21.33 | 21.19 | 0.89 | 21.62 | 21.05 | 0.61 |
|  | (3.96) | (4.27) | (3.74) |  | (4.20) | (3.86) |  |
| BIS Non- | 13.22 | 13.04 | 13.39 | 0.62 | 12.57 | 13.59 | 0.16 |
| planning | (2.63) | (2.68) | (2.62) |  | (2.20) | (2.80) |  |
| Center for | 8.53 | 8.85 | 8.26 | 0.64 | 7.81 | 8.95 | 0.39 |
| Epidemiologic | (4.76) | (5.44) | (4.15) |  | (3.76) | (5.24) |  |
| Studies |  |  |  |  |  |  |  |
| Depression |  |  |  |  |  |  |  |
| Scale |  |  |  |  |  |  |  |
| Snaith-Hamilton | 1.14 | 1.33 | 0.97 | 0.59 | 1.05 | 1.19 | 0.84 |
| Pleasure Scale | (2.54) | (2.99) | (2.12) |  | (3.04) | (2.26) |  |
| PANAS (+) | 33.28 | 31.15 | 35.13 | 0.12 | 33.90 | 32.92 | 0.71 |
|  | (9.59) | (10.0 | (8.89) |  | (8.74) | (10.14) |  |
|  |  | 8) |  |  |  |  |  |
| PANAS (-) | 17.62 | 18.19 | 17.13 | 0.60 | 16.95 | 18.00 | 0.62 |
|  | (7.64) | (7.24) | (8.05) |  | (8.37) | (7.28) |  |
| Power of Food | 51.88 | 48.70 | 54.65 | 0.24 | 50.00 | 52.95 | 0.58 |
| Scale | (19.22) | (19.8 | (18.5 |  | (21.7) | (17.87) |  |
|  |  | 8) | 0) |  |  |  |  |
| Food Cravings | 101.69 | 99.07 | 103.9 | 0.62 | 96.62 | 104.57 | 0.43 |
| Questionnaire | (36.56) | (37.6 | 7 |  | (37.19) | (36.40) |  |
|  |  | 5) | (36.0 |  |  |  |  |
|  |  |  | 5) |  |  |  |  |
| WREQ routine | 1.64 | 1.43 | 1.83 | 0.08 | 1.46 | 1.75 | 0.23 |
| restraint | (0.87) | (0.58) | (1.04) |  | (0.69) | (0.95) |  |

|  |  |  |  |  |  |  |  |  |
| --- | --- | --- | --- | --- | --- | --- | --- | --- |
| WREQ | 2.17 | 2.05 | 2.27 | 0.32 | 2.17 | 2.16 | 0.96 | 616 |
| compensatory | (0.84) | (0.81) | (0.85) |  | (0.92) | (0.80) |  |  |
| restraint |  |  |  |  |  |  |  |  |
| WREQ | 2.30 | 2.21 | 2.37 | 0.57 | 2.41 | 2.23 | 0.52 |  |
| susceptibility to | (1.00) | (1.06) | (0.96) |  | (1.15) | (0.92) |  |  |
| external cues |  |  |  |  |  |  |  |  |
| WREQ | 2.09 | 1.95 | 2.21 | 0.34 | 2.13 | 2.06 | 0.79 |  |
| emotional | (1.02) | (1.18) | (0.86) |  | (1.15) | (0.95) |  |  |
| eating |  |  |  |  |  |  |  |  |
| SLIM | 0.72 | -3.24 | 4.30 | 0.46 | -11.70 | 8.11 | 0.06 |  |
|  | (39.15) | (33.4 | (43.9 |  | (36.43) | (39.31) |  |  |
|  |  | 5) | 2) |  |  |  |  |  |
| Number of | 11 | 14 | 8 | 0.21 | 14 | 9 | 0.22 |  |
| candies eaten | (17.19) | (20.0 | (13.9 |  | (19.66) | (15.42) |  |  |
|  |  | 2) | 4) |  |  |  |  |  |

**Table 2.** Demographic, biometric, and self-reported data for crossed participant groups

| Characteristic | Mean (SD) |  |  | Mean (SD) |  | Mean (SD) |  |
| --- | --- | --- | --- | --- | --- | --- | --- |
|  | Group 00 | Group 01 | *p* (00 | Group 10 | *p* (00 | Group 11 | *p* (00 |
|  | (*n* = 20) | (*n* = 11) | vs. 01) | (*n* = 17) | vs. 10) | (*n* = 11) | vs. 11) |
| Age, years | 47 (9.15) | 48 (7.62) | 0.77 | 46 (14.26) | 0.76 | 37<br>(12.63) | 0.02 |
| Women, % | 50 | 55 | 0.81 | 47 | 0.86 | 27 | 0.22 |
| Race, % |  |  |  |  |  |  |  |
| Black/African American | 50 | 64 |  | 71 |  | 82 |  |
| White/Caucasian | 35 | 27 |  | 18 |  | 9 |  |
| Asian | 5 | 0 |  | 12 |  | 9 |  |
| More than one race | 5 | 9 |  | 0 |  | 0 |  |
| I prefer not to say | 5 | 0 |  | 0 |  | 0% |  |
| BMI, kg/m<sup>2</sup> | 30 (5.73) | 33<br>(9.83) | 0.28 | 32<br>(9.11) | 0.46 | 28 (6.15) | 0.34 |
| Hispanic or Latino ethnicity, % | 10 | 27 |  | 12 |  | 0 |  |
| BIS attention | 15.20<br>(3.38) | 16.36<br>(2.94) | 0.35 | 15.94 (5.08) | 0.60 | 14.80 (1.32) | 0.72 |
| BIS motor | 21.20<br>(3.07) | 21.18<br>(4.90) | 0.99 | 20.88 (4.72) | 0.81 | 22.10 (3.48) | 0.47 |
| BIS Non-planning | 13.65<br>(2.85) | 12.91 (2.17) | 0.46 | 13.53 (2.83) | 0.90 | 12.20 (2.30) | 0.17 |
| Center for<br>Epidemiologic<br>Studies<br>Depression Scale | 8.05 (4.02) | 8.64 (4.54) | 0.71 | 10.00 (6.36) | 0.27 | 6.90 (2.60) | 0.42 |
| Snaith-Hamilton<br>Pleasure Scale | 1.45 (2.52) | 0.09 (0.30) | 0.09 | 0.88 (1.93) | 0.45 | 2.10 (4.25) | 0.60 |
| PANAS (+) | 34.95<br>(9.60) | 35.45 (7.85) | 0.88 | 30.53 (10.52) | 0.19 | 32.20 (9.75) | 0.47 |
| PANAS (-) | 16.25<br>(6.26) | 18.73<br>(10.75) | 0.42 | 20.06 (8.02) | 0.11 | 15.00 (4.37) | 0.58 |
| Power of Food<br>Scale | 54.45<br>(17.61) | 55.00<br>(20.90) | 0.94 | 51.18 (18.56) | 0.59 | 44.50 (22.32) | 0.19 |
| Food Cravings<br>Questionnaire | 105.05<br>(37.51) | 102.00<br>(34.93) | 0.83 | 104.00<br>(36.19) | 0.93 | 90.70 (40.54) | 0.34 |
| WREQ routine<br>restraint | 1.98 (1.13) | 1.55 (0.82) | 0.27 | 1.47 (0.61) | 0.10 | 1.37 (0.53) | 0.12 |
| WREQ<br>compensatory<br>restraint | 2.28 (0.81) | 2.24 (0.97) | 0.90 | 2.02 (0.79) | 0.32 | 2.10 (0.90) | 0.58 |
| WREQ<br>susceptibility to<br>external cues | 2.25 (0.83) | 2.58<br>(1.18) | 0.37 | 2.21 (1.05) | 0.90 | 2.22 (1.14) | 0.94 |
| WREQ emotional<br>eating | 2.12 (0.73) | 2.36 (1.07) | 0.46 | 1.99 (1.18) | 0.68 | 1.88 (1.24) | 0.51 |
| SLIM | 17.95<br>(39.16) | -20.50<br>(42.70) | 0.02 | -3.45 (37.31) | 0.10 | -2.91 (28.17) | 0.13 |
| Number of candies eaten | 4 (8.89) | 15 (18.76) | 0.04 | 14 (19.74) | 0.06 | 14 (21.42) | 0.09 |

### 2.2. Study procedures

The study consisted of an eligibility screening of potential participants via telephone followed by an in-person laboratory visit. A research assistant met with the participant at each laboratory visit to explain the study and obtain informed consent. The research assistant then collected the participant’s biometric information, including height and weight, and then administered a series of computerized questionnaires. After completing the questionnaires, the research assistant placed an EEG net on the participant’s head and instructed the participant on how to complete the cued food delivery task. The research assistant then left the room and began both the EEG recording and the cued food delivery task. After completion of the EEG session and task, the participant was debriefed and given financial compensation. All study procedures were approved by The University of Texas MD Anderson Cancer Center Institutional Review Board.

### 2.3. Questionnaires

Our computerized questionnaires consisted of those assessing hunger and satiety, eating habits, impulsivity, mood, affect, and hedonic tone. To assess hunger and satiety, we administered the Satiety Labeled Intensity Magnitude (SLIM) scale [29] to each participant before and after completion of the cued food delivery task. To ascertain eating habits, we used the weight-related eating questionnaire (WREQ) [30], the Power of Food Scale [31], and the Food Cravings Questionnaire [32], which measure variables such as susceptibility to external cues, the influence of a food-abundant environment on eating, and food cravings, respectively. To measure impulsivity, we administered the Barratt Impulsiveness Scale (BIS) [33]. Finally, to identify variables relating to affect, mood, and hedonic tone, we administered the Positive and Negative Affect Schedule (PANAS) [34], the Center for Epidemiologic Studies Depression Scale [35], and the Snaith-Hamilton Pleasure Scale [36], respectively.

### 2.4. Cued food delivery task

Participants completed the cued food delivery task depicted in Fig. 1 [18] with the addition of a control condition in which they were also dispensed plastic beads. During the task, participants viewed emotional, neutral, and food-related images presented on a 17-inch computer screen using E-Prime software (version 2.0.8.74; Psychology Software Tools, Inc., Pittsburgh, PA) while EEG was recorded from the scalp. After viewing a food-related image, each participant was dispensed a chocolate candy, which they had the option to eat or discard, or a bead. Food-related images consisted salty or sweet contents (for example: pizza [salty], cake [sweet]). One of these two categories of food images (counterbalanced across participants) preceded the delivery of the candy, whereas the other preceded the delivery of the bead. Each participant was told at the beginning of the EEG session which category of food image would precede the candies and which would precede the beads. The pictures used in this task were selected from the International Affective Picture System (IAPS) [37] and a set of pictures used in our previous studies [15,16]. See Supplementary Table 1 for a list of the IAPS pictures used in the experiment.

**Fig. 1.**
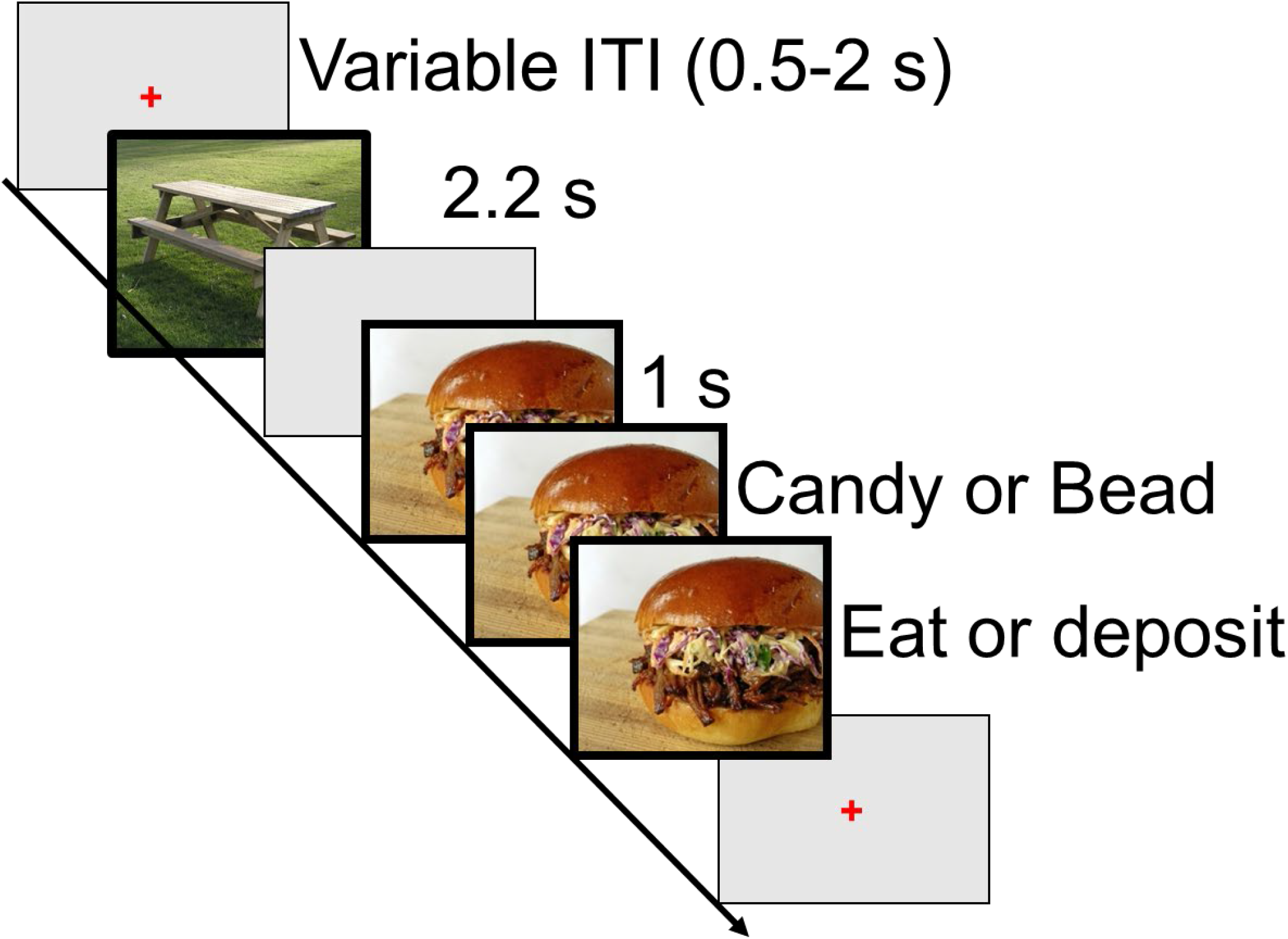
A schematized representation of the timing of events in the cued food delivery task. In this task, participants viewed emotional, neutral, and food-related images and made food-related decisions. After a food-related image was presented on the screen, the participant received either a candy (reward condition), which they could eat or discard, or a bead (control condition). ITI: intertrial interval.

The cued food delivery task consisted of six experimental blocks that lasted about 5 min each. In each block, 55 images were presented pseudorandomly (no more than two images belonging to the same picture category were presented consecutively): 10 neutral (people and objects), 10 pleasant (erotica and romance), 15 unpleasant (mutilations, violence, and pollution), and 20 food-related (savory or sweet) images.

No images were repeated during the task. For the food images, the candy or bead was dispensed 1000 ms after the food cue appeared on the screen through a tube into a receptacle. The participant then could either pick up and eat the candy or discard it in a box. Each food image remained visible until the participant either deposited the candy or bead into the box or pressed a button indicating that they had finished eating the candy. All non-food images were presented for 2.2 s, and a random intertrial interval (ITI) of 3-5 s separated each trial. To familiarize the participants with the task, we ran 11 practice trials, two of which were followed by a candy or bead.

### 2.5. EEG acquisition

We continuously recorded EEG during the task using a 129-channel Geodesic Sensor Net that was amplified with an AC-coupled high-input-impedance (200 MΩ) amplifier (Geodesic EEG System 200; EGI, Eugene, OR) and referenced to electrode Cz. EEG data were collected at a sampling rate of 250 Hz and filtered online using a 0.1-Hz high-pass and 100-Hz low-pass filter. The scalp impedance was kept under 50 KΩ as per the manufacturer’s instructions.

### 2.6. Data reduction

After collecting the EEG, the EEG data were filtered using a 30-Hz low-pass filter and visually inspected to identify broken channels, which were defined as any channels contaminated by artifacts in more than 50% of the recording. Any broken channels were interpolated using spherical splines. Next, the EEG recordings were corrected for blinks and horizontal eye movements using a spatial filtering method implemented in the BESA software program (version 5.1.8.10; MEGIS Software GmbH, Gräfelfing, Germany). Next, the data were transformed to the average reference and segmented as follows. For the analysis of ERPs, each segment of EEG was time-locked to the onset of each picture in segments that started 1500 ms before the onset of the picture and lasted until 1500 ms afterward. For the time-frequency analyses, each segment of EEG was time-locked to the delivery of a candy or bead in segments that started 1500 ms before the dispensation of the candy or bead and lasted until 1500 ms afterward. The data were baseline-corrected using a 100-ms time bin before the onset of the pictures (ERPs) or the onset of the candy or bead dispensation (time-frequency) as the baseline. Artifacts in the −1000- to +1000-ms time window for each segment were then detected based on the following criteria: EEG amplitude above 100 or below −100 μV, an absolute voltage difference between any two points in a segment no greater than 100 μV, maximum voltage step between two contiguous data points of 20 μV, and less than 0.5 μV of variation in activity for more than 100 ms. Channels that were marked bad in more than 40% of the segments were interpolated, and any segment that included more than 12 bad channels after interpolation was discarded.

### 2.7. LPP

We used the amplitude of the LPP as a measure of cues’ motivational salience. To calculate the LPP for each subject and picture category, the EEG responses that were time-locked to the onset of each picture during the 400- to 800-ms time window using a pooled set of centroparietal sensors (EGI HydroCel Geodesic Sensor Net sensors 7, 31, 37, 54, 55, 79, 80, 87, 106, 129; see also Fig. 2 inset) were averaged together. This is the same temporospatial region of interest (ROI) used in our previous studies investigating the LPP [15,38,39].

**Fig. 2.**
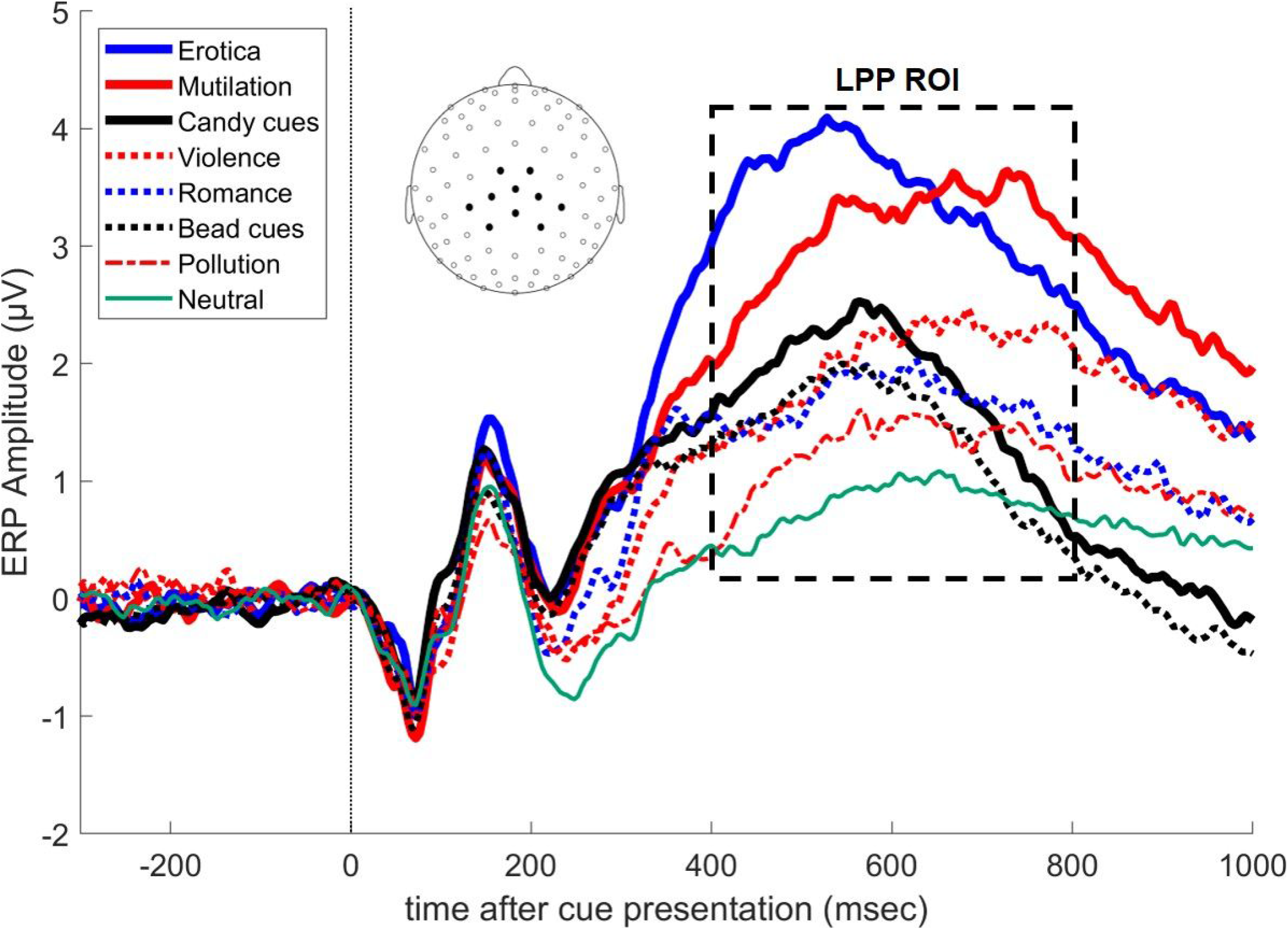
ERPs for centroparietal scalp sites (see inset for EEG electrode locations) showing that, on average, the LPP amplitude was higher for motivationally relevant pictures, such as mutilations or erotic images, than for other types of pictures. The box outlines the temporal ROI used to calculate the LPP for each picture category. Each picture was presented on the screen at time 0.

### 2.8. Theta power

To calculate theta power, the EEG data time-locked to the delivery of the candy or bead were transformed into the time-frequency domain using a continuous wavelet transform. The wavelet transform was based on a complex Morlet wavelet function with a Morlet parameter of 5 using 40 linear frequency steps from 1 to 40 Hz. The data were normalized using Gabor normalization and were baseline-corrected using a reference interval from −875 to −625 ms. To calculate theta power, the 4- to 8-Hz frequency bands were averaged.

### 2.9. Classification of participants

To classify the participants based on their LPP responses to cues, we followed a procedure used in our previous studies [15,16]. Specifically, we *z*-transformed each participant’s LPP data for each picture category, then we applied a *k*-means (*k* = 2) clustering algorithm to these *z*-transformed LPP values for each participant. The number of clusters (*k* = 2) was decided a priori based on previous findings [16,38].

To classify the participants based on their theta power amplitudes, we followed a similar strategy Specifically, each participant’s theta power values were *z*-transformed for the candy, bead, and neutral conditions during a 0- to 200-ms time bin using a pooled set of mid-frontal sensors. We then applied a *k*-means (*k* = 2) clustering algorithm to these z-transformed theta power values, with the a priori hypothesis that two distinct patterns of theta activity would be observed, much like our previous findings using the LPP.

### 2.10. Eating behavior

Because the number of candies the participants ate during the experiment is a count variable, we tested differences in eating behavior between participant groups using Poisson regression analysis. First, we compared the number of candies eaten during the experiment between the two LPP-derived groups. Second, we conducted another Poisson analysis to compare the number of candies eaten between the two theta power-derived groups. Third, we compared the number of candies eaten by the four groups formed by crossing the LPP and theta power-based groups using Poisson regression.

### 2.11. Demographics and questionnaires

To identify whether any demographic or psychological factors had confounding effects on the trends in eating behavior we observed in the participant groups, we conducted Poisson regression modeling the effect of group assignment (LPP and theta power) on eating behavior. The demographic, biometric, and self-reported data outlined in Table 1 were included in the model as covariates.

## 3. Results

### 3.1. ERPs

Fig. 2 shows the grand averaged ERPs for each picture category. As expected, the amplitude of the LPP increased as a function of motivational salience irrespective of hedonic content. We formally tested this effect using LPP amplitude as a dependent variable in a repeated-measures analysis of variance (ANOVA) with the picture category as an eight-level factor (candy cues, bead cues, erotica, romance, neutral, pollution, violence, and mutilations; *F*[7, 399] = 22.1, *p* < 0.001). We also tested the quadratic trend of increasing LPP as a function of motivational salience under both pleasant and unpleasant conditions (*F*[5, 290] = 46.3, *p* < 0.001). Furthermore, we found that on average, food images preceding dispensation of the candy elicited larger LPPs than did food images preceding dispensation of the bead (*F*[1, 58] = 5.02, *p* = 0.029).

### 3.2. Classification of participants: LPP

Cluster analysis of the LPP responses identified the two hypothesized reactivity profiles: one group (C>P) had larger LPP responses to food cues than to pleasant images, and the other group (P>C) had larger LPP responses to pleasant images than to food cues. Both groups exhibited the canonical pattern of progressively larger LPP responses for both pleasant and unpleasant images as a function of their motivational salience (C>P group: *F* = 87.5, *p* < 0.001; P>C group: *F* = 77, *p* < 0.001) (Fig. 3A). However, the P>C group had significantly larger LPP responses to pleasant images than to food cues (*F* = 6.51, *p* = 0.013), whereas the C>P group had significantly larger LPP responses to food cues than to pleasant images (*F* = 59, *p* < 0.001). See Supplementary Fig. S1 for the averaged LPP amplitudes across all pleasant picture contents by group assignment.

**Fig. 3.**
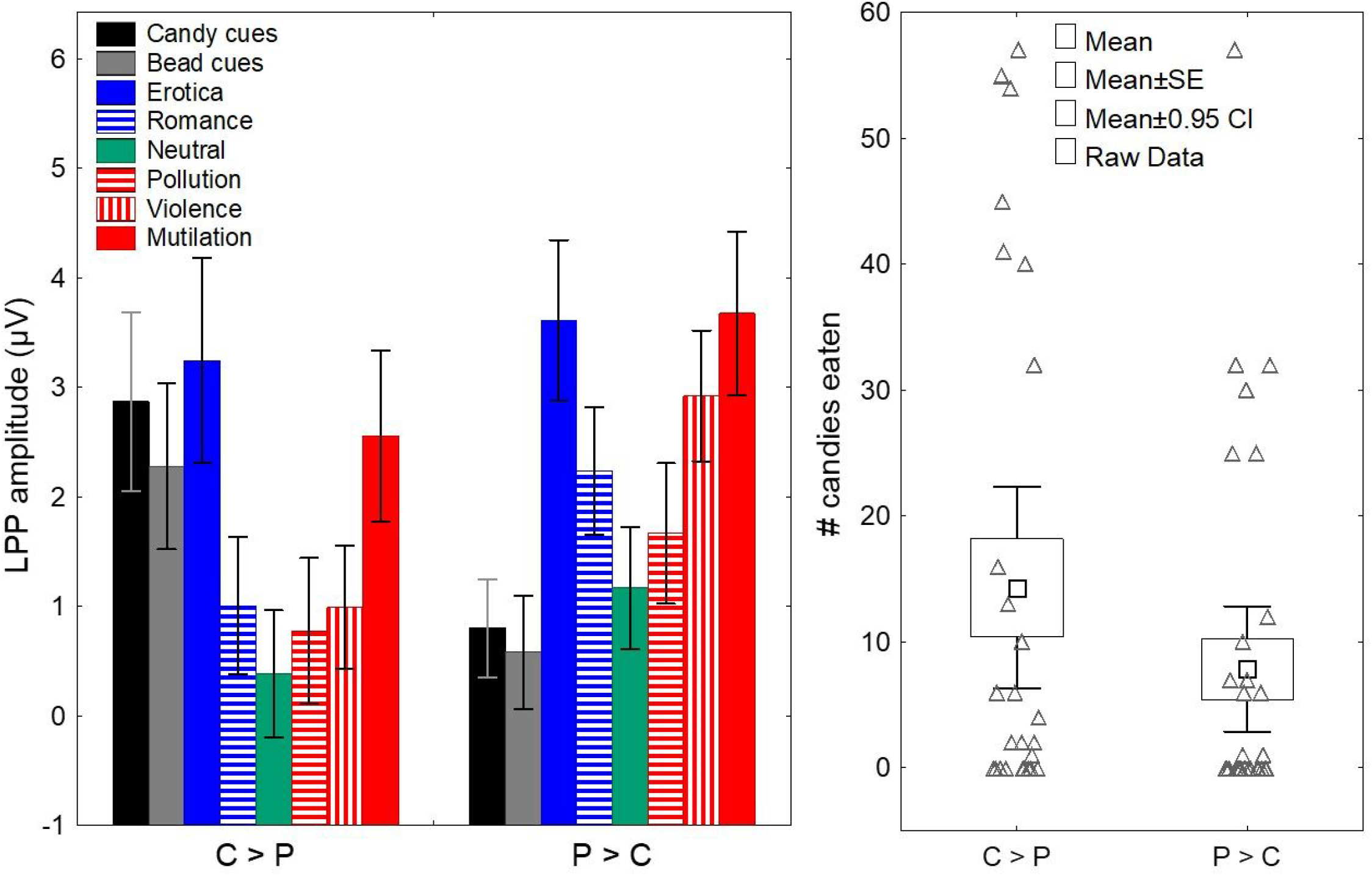
We replicated the finding that individual differences in LPP responses to food cues & non-food-related pleasant images are predictive of cue-induced eating. (**A**) *K*-means clustering of the LPP responses identified two groups: one with higher LPP amplitudes for food cues than for pleasant images (C>P group) and one with higher LPP amplitudes for pleasant images than for food cues (P>C group). Both groups exhibited the canonical pattern of progressively larger LPP responses for both pleasant and unpleasant images as a function of their motivational salience. Error bars: 95% CI. (**B**) The C>P group ate significantly more candies during the cued food delivery task than did the P>C group. SE: standard error. Error bars: 95% CIs.

After determining the group assignment for each participant, we then compared the number of candies eaten by the C>P and P>C groups (Fig. 3B). Poisson regression analysis demonstrated that individuals in the C>P group ate significantly more during the experiment than did individuals in the P>C group (Wald X^2^[1] = 43.1, p<0.001). Demographic, biometric, and self-reported questionnaire data for the C>P and P>C groups are reported in Table 1.

### 3.3. Time-frequency power

To identify a set of EEG sensors to pool together in our analysis of theta power, we used the following procedure. Using theta power as a dependent variable, we performed a repeated-measures ANOVA with condition (candy, bead, and neutral) as a factor for each time point and each EEG sensor. To identify the sensors and time points at which theta power exhibited statistically significant differences across conditions, we determined thresholds for the *F*-values resulting from these ANOVAs using Bonferroni correction. We then selected the sensors that showed statistically significant differences between conditions (candy, bead & neutral) during the 0- to 200-ms time bin (see inset in Fig. 4 for this set of sensors), during which cognitive control-related effects in theta power are typically greatest [25]. See Supplementary Fig. S2 for the topography of these *F*-values during this time bin. We then averaged theta power in the 0- to 200-ms time bin from this pooled set of sensors to obtain a single theta power value for each participant under the candy, bead, and neutral conditions. Figure 4 shows the time course of theta power in the Candy, Bead & Neutral conditions. We found that, on average, power increased when either candies or beads were dispensed to the participant, but not when they passively viewed neutral pictures.

**Fig. 4.**
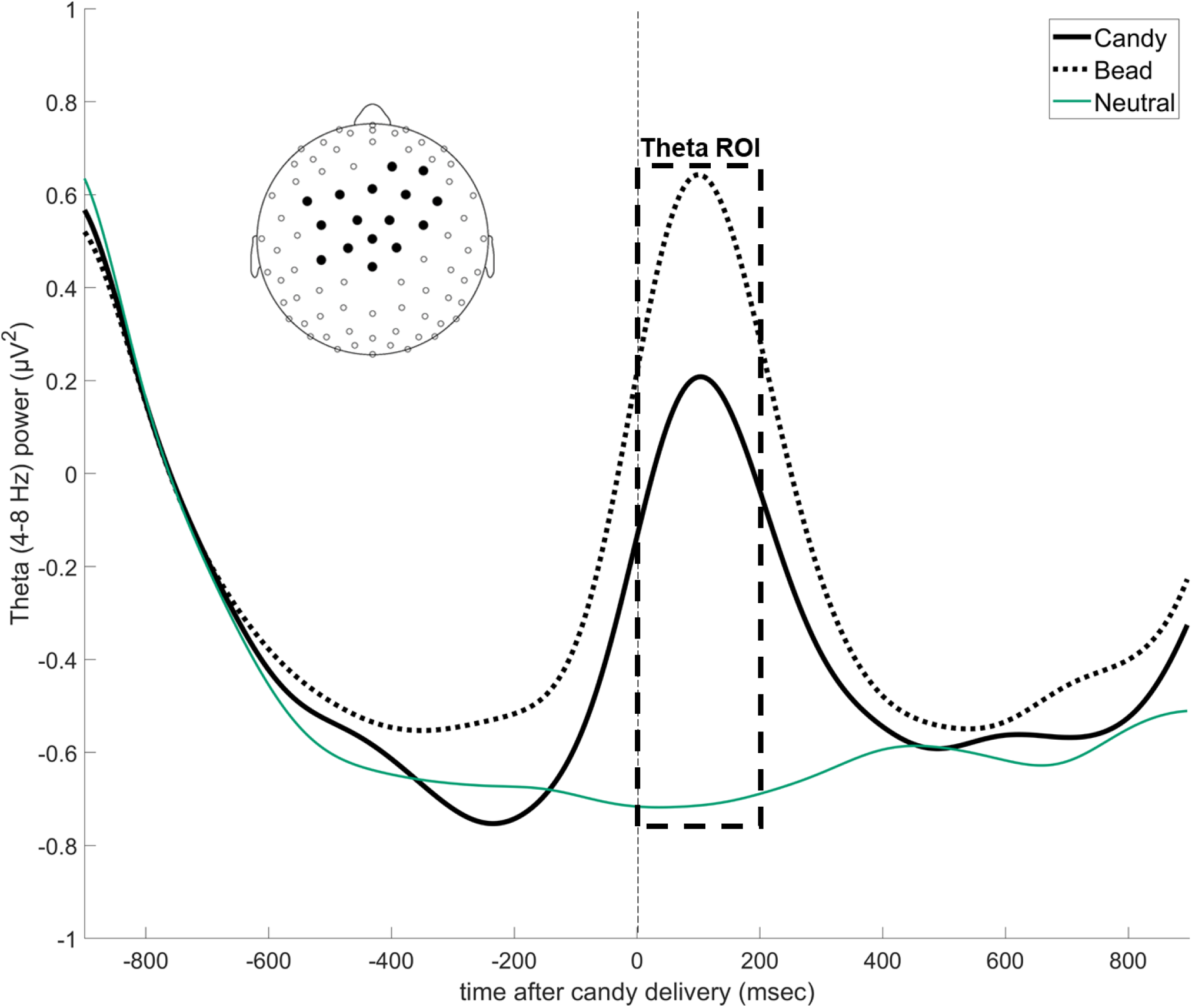
Time series data showing changes in average theta power over mid-frontal scalp sites during the candy, bead & neutral experimental conditions. Theta power over midfrontal scalp sites (see inset for EEG electrode locations) increases during the candy and bead conditions but not when the participant is passively viewing neutral pictures. The box indicates the temporal ROI used to calculate theta power. The candies and beads were delivered at time 0.

Next, to determine whether theta power differed between the C>P and P>C groups, we averaged the pooled and binned theta power values for these two groups. We found that the groups had similar dynamics in theta power under the candy, bead, and neutral conditions. A repeated-measures ANOVA demonstrated no significant interaction effect of group assignment (C>P and P>C) and condition (candy, bead, and neutral) (*F*[2, 114] = 0.667, *p* = 0.515) on theta power. These data are shown in Supplementary Fig. S3.

### 3.4. Classification of participants: theta power

Cluster analysis of theta power identified two participant groups (Fig. 5A): one with higher theta power for the candy condition than for the bead condition (θCA>θBE group) and the other with higher theta power for the bead condition than for the candy condition (θBE>θCA group). We then compared the number of candies eaten by these two groups during the experiment (Fig. 5B). Poisson regression analysis demonstrated that the θCA>θBE group ate significantly more candies than did the θBE>θCA group (Wald *X*^2^=[1] = 41.5, *p* < 0.001). Demographic, biometric, and self-reported questionnaire data for the θCA>θBE and θBE>θCA groups are reported in Table 1.

**Fig. 5.**
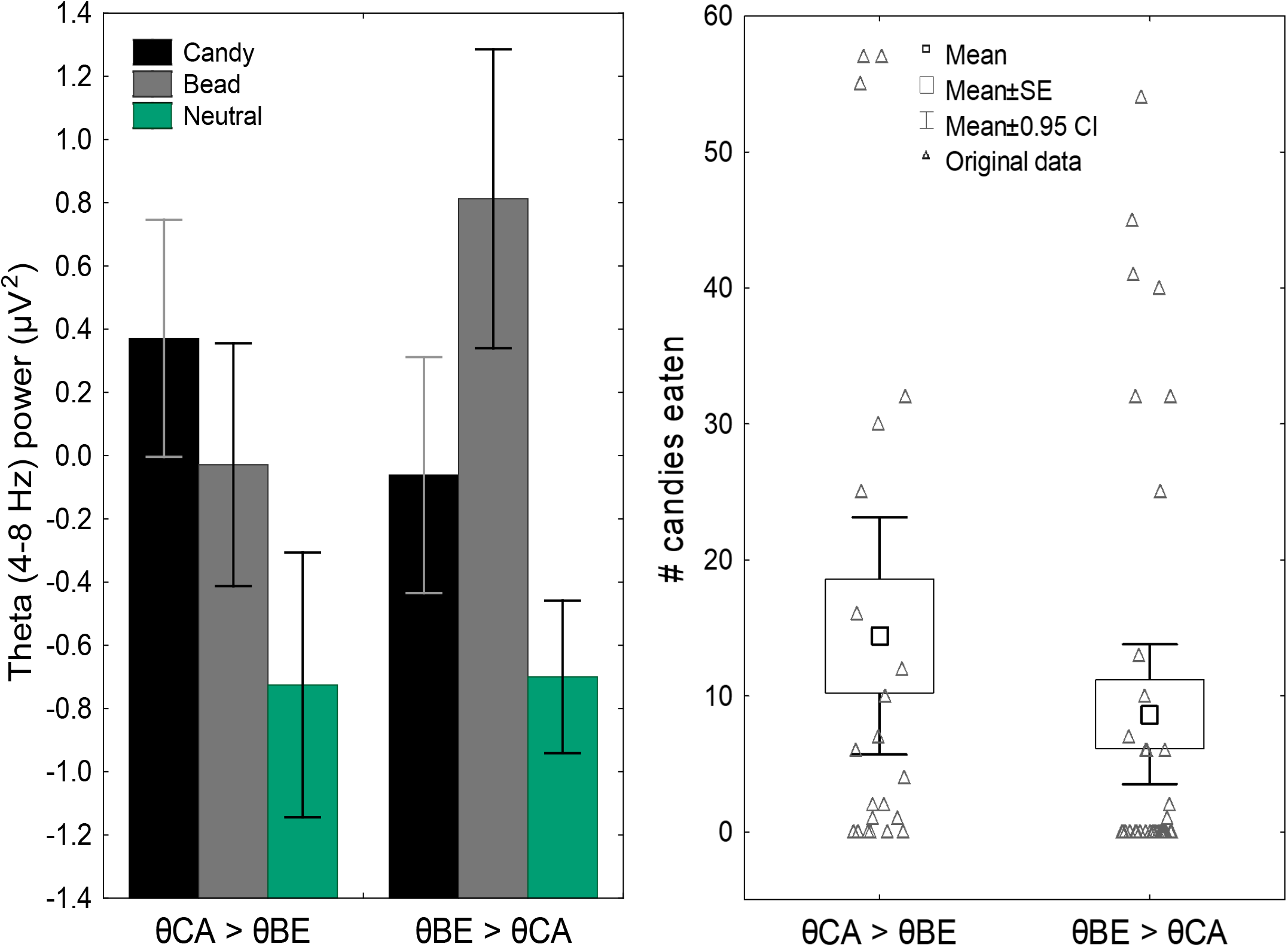
We found that individual differences in theta power during food-related decision-making were predictive of cue-induced eating. (**A**) *K*-means clustering of theta power data identified two groups: one with higher theta power for the candy condition than for the bead condition (θCA>θBE group) and one with higher theta power for the bead condition than for the candy condition (θBE>θCA group). Error bars: 95% CIs. (**B**) The θCA>θBE group ate significantly more candies during the cued food delivery task than did the θBE>θCA group. SE: standard error. Error bars: 95% CI.

### 3.5. Classification of participants: LPP and theta power

Next, to determine how both individual differences in the attribution of motivational salience to food cues and the engagement of cognitive control confer vulnerability to cue-induced eating, we created four participant groups by crossing the results of the LPP and theta power classification procedures. We labeled these four groups 00 (the P>C and θBE>θCA groups), 01 (the P>C and θCA>θBE groups), 10 (the C>P and θBE>θCA groups), and 11 (the C>P and θCA>θBE groups). Demographic, biometric, and self-reported questionnaire data for these four crossed groups are reported in Table 2.

After crossing the group assignments for both the LPP and theta power cluster analyses, Poisson regression analysis demonstrated a significant effect of group assignment on the number of candies eaten during the experiment (Wald *X*^2^=[3] = 106.2, *p* < 0.001). Notably, although the individuals in group 00 ate the least, those in the three remaining groups had similar levels of eating behavior on average (Wald *X*^2^=[2] = 0.825, *p* = 0.662) (Fig. 6).

**Fig. 6.**
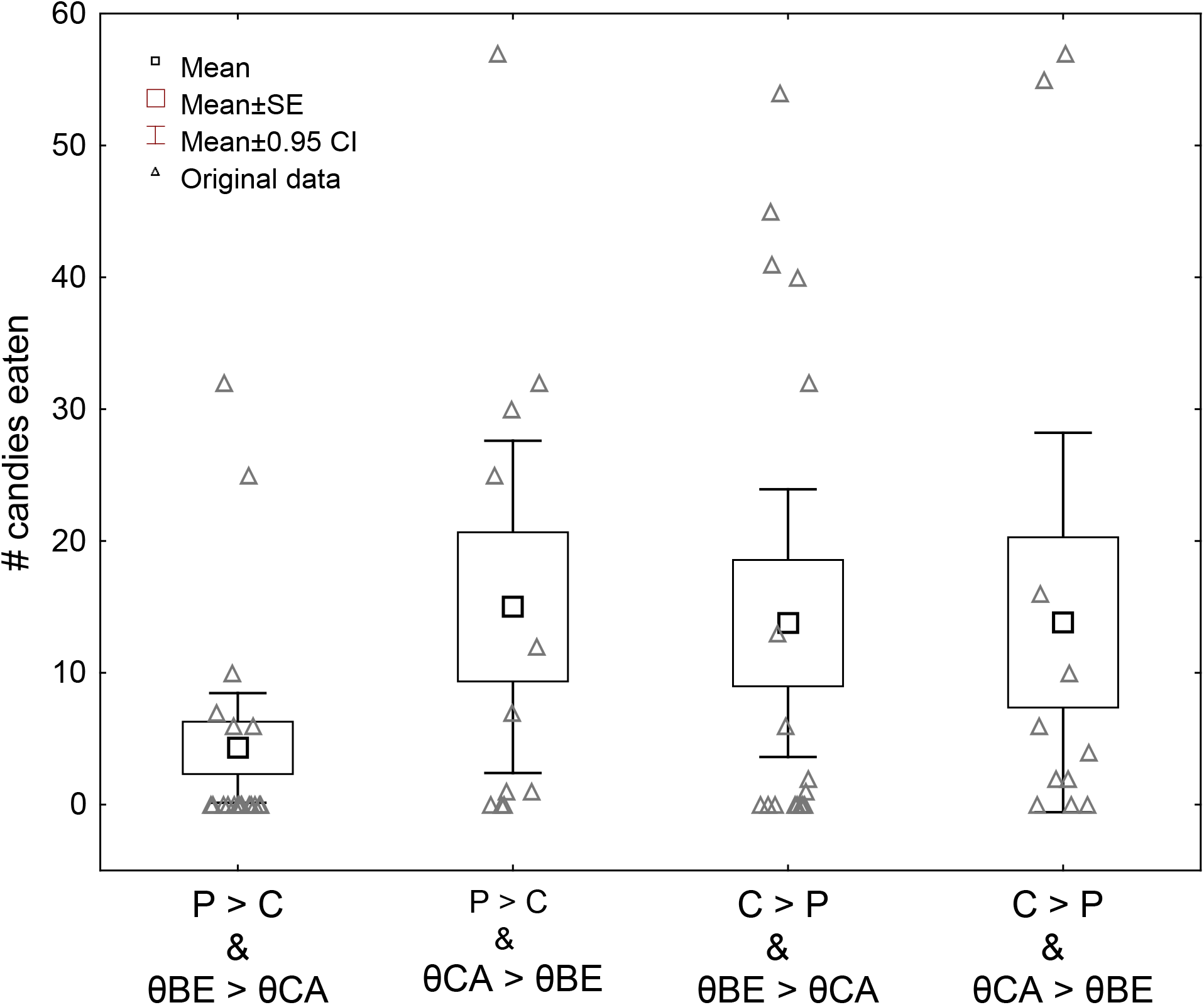
Number of candies eaten per group, by crossed LPP and theta power groups. After crossing the participant groups according to the LPP and theta-based cluster analyses, we found that individuals with neither LPP nor theta-based “risk factors” (P>C and θBE>θCA [group 00]) ate the least of all four groups. The three remaining groups exhibited similar levels of eating behavior on average. SE: standard error. Error bars: 95% CIs.

### 3.6. Demographics and covariates

We conducted Poisson regression analysis modeling the effect of crossed group assignment on eating behavior which included the demographic, biometric, and questionnaire data outlined in Table 2 as covariates. We found a significant main effect of group assignment on eating behavior for all groups except for those with both LPP and theta risk factors (group 01 AKA the P>C and θCA>θBE group) after controlling for factors such as hunger and satiety [29], eating behavior [31,32], sensitivity to reward and punishment [40], mood [34], and impulsivity [33].

## 4. Discussion

This study aimed at determining the role that individual differences in affective and cognitive brain systems have in regulating cue-induced eating. This work is informed by results from animal models demonstrating that individual differences in top-down attentional control and bottom-up attribution of motivational salience to food-related cues influence reward-seeking behaviors. By investigating both food-related decision-making and the motivational salience of cues, we aimed to elucidate how these mechanisms contribute to cue-induced eating behavior in humans. Because we found that both LPP and theta power-based groups showed statistically significant differences in eating behavior, we may conclude that these mechanisms likely act independently to regulate eating behavior during the cued food delivery task

By applying cluster analysis to the LPP responses evoked by food-related and non-food-related motivationally salient images, we identified two reactivity profiles associated with vulnerability to cue-induced eating: individuals with larger LPP responses to food-related cues than to pleasant images (C>P group) ate significantly more than did individuals with larger LPP responses to pleasant stimuli than to food-related cues (P>C group). These results replicate those from previous studies [15,16] and support the hypothesis that individual differences in the tendency to attribute motivational salience to food-related cues than to other pleasant stimuli underlie vulnerability to cue-induced eating [12,13,39,41].

Furthermore, we found that midfrontal theta power increased after the delivery of candies & beads and that individual differences in midfrontal theta power were associated with vulnerability to cue-induced eating. Specifically, individuals with higher phasic theta power following the delivery of candies than of beads (θCA>θBE) ate more during the cued food delivery task than did individuals with the opposite theta response pattern (θBE>θCA). Authors have proposed that changes in theta power over midfrontal scalp sites represent an index of the engagement of cognitive control mechanisms [25] because midfrontal theta tends to increase when an individual is executing a task that requires increased attentional demands, such as inhibiting prepotent responses [26,27] and performing otherwise cognitively demanding tasks [28]. In light of these findings, the results of the present study suggest that some individuals struggle with food-related decision-making and that these individuals are more likely to engage in cue-induced eating when a palatable food option is available [10,42].

In addition, we found that the LPP-based and θ-based reactivity profiles reflect affective and cognitive mechanisms that independently contribute to cue-induced eating. This is evidenced by the finding that the C>P and P>C groups had similar theta power dynamics during food-related decision-making. Furthermore, after crossing the group assignments for the LPP and theta-based cluster analyses, we found that the group with neither the LPP nor the theta risk factor (P>C and θBE>θCA group) ate the least of all four groups and that the three remaining groups exhibited similar levels of eating behavior on average. These results imply that individuals at risk for cue-induced behaviors due to the presence of both LPP and theta-based risk factors are no more vulnerable to cue-induced behaviors than are those who have only one of these two risk factors. Further studies are needed to determine if this finding is consistent across populations and paradigms.

Whereas the validity of the LPP in predicting cue-induced behavior has been well replicated [16,38,43] and is consistent with theoretical models concerning the motivational salience of cues [12,14], the predictive validity of a theta-based correlate described in the present study is novel and should be considered preliminary until it is replicated. Although in previous studies researchers have used theta power to index the engagement of higher cognitive functions during the execution of cognitively demanding tasks [25], in the present study, we did not explicitly manipulate cognitive load during food-related decision-making. Because we did not explicitly manipulate cognitive control via a cognitively demanding task, inferring from our results that the observed dynamics in theta power are in fact due to the engagement of higher cognitive functions remains speculative.

Also, although our results suggest that individual differences in the ability to exert cognitive control are independent from the tendency to attribute motivational salience to cues, studies using preclinical models suggested that these two cognitive-motivational styles are coupled: animals with high motivational salience attributed to food-related cues are also more impulsive and less able to implement top-down attentional control in the presence of cues than are those who do not attribute high motivational silence to food-related cues [12,44–46]. In light of these incongruous findings, theta power analysis as implemented in the present study may not capture the same aspects of cognitive control that are captured using animal models, which may be more related to impulsivity than to top-down attentional control. Our self-reported data did not demonstrate significant differences in impulsivity scores between groups, which may explain the divergent findings of the present study and those in the animal literature. Meanwhile, results from human and animal studies of cue-induced behavior may be inconsistent because of the inherent differences between humans and animal models [41]: complex human behaviors result from a more evolved cognitive control system [10] than that of animal models and thus might not be probed as effectively using animal behavioral approaches. Further studies leveraging the paradigm that we used here will facilitate the bidirectional translation and improvement of both animal models and human subjects research investigating cue-induced behavior.

Our findings are consistent with those for neurobiological models suggesting that both high reactivity to food-related cues and deficits in cognitive control can lead to excessive eating [11]. Furthermore, because our results suggest that cognitive control and reward networks independently contribute to cue-induced eating, these findings further emphasize the need for individualized treatments of maladaptive reward-seeking behaviors. This has worthwhile clinical implications: by separating the roles of both affective and cognitive psychophysiological correlates in predicting cue-included eating, we may identify potential biomarkers of vulnerability to overeating and obesity that could guide treatment decisions. For example, using repetitive transcranial magnetic stimulation (rTMS), a non-invasive neuromodulation technique [47], we can upregulate brain activity in cognitive control networks [48] or downregulate brain activity in reward networks [49]. Thus, a patient identified to have high affective vulnerability may be selected for inhibitory rTMS of the ventromedial prefrontal cortex, which is commonly implicated in reward processing [50]. Also, a patient with cognitive vulnerability may be more effectively treated with excitatory rTMS of the dorsolateral prefrontal cortex, which is commonly implicated in executive control [51].

In conclusion, our results demonstrated that both the amplitude of the LPP and theta power are predictive of cue-induced eating behavior, suggesting that both affective and cognitive mechanisms are implicated in the regulation of cue-induced eating. By simultaneously measuring both the amplitude of the LPP and theta power while participants were in the presence of food-related cues and actual food rewards, we clarified the mechanisms underlying cue-induced eating behaviors. Continuing this line of investigation may inform clinicians of mechanisms underlying maladaptive eating and may foster the development of personalized clinical interventions for excessive eating.

## Abbreviations

ANOVA: analysis of variance
BIS: Barratt Impulsiveness Scale
C>P: LPP responses that are larger to food cues than to pleasant images
EEG: electroencephalogram
ERP: event-related potential
LPP: late positive potential
P>C: LPP responses that are larger to pleasant images than to food cues
PANAS: Positive and Negative Affect Scale
ROI: region of interest
SLIM: Satiety Labeled Intensity Magnitude
θBE>θCA: theta power that is higher under the bead condition than under the candy condition
θCA>θBE: theta power that is higher under the candy condition than under the bead condition
WREQ: weight-related eating questionnaire

## Acknowledgment

We thank Drs. Scott Lane and Andreas Keil for advising us in the analysis of these data and MD Anderson Editing Services, Research Medical Library for assistance in editing the manuscript.

## Supplementary Materials

### Supplementary Figure Captions

**Supplementary Table 1.**
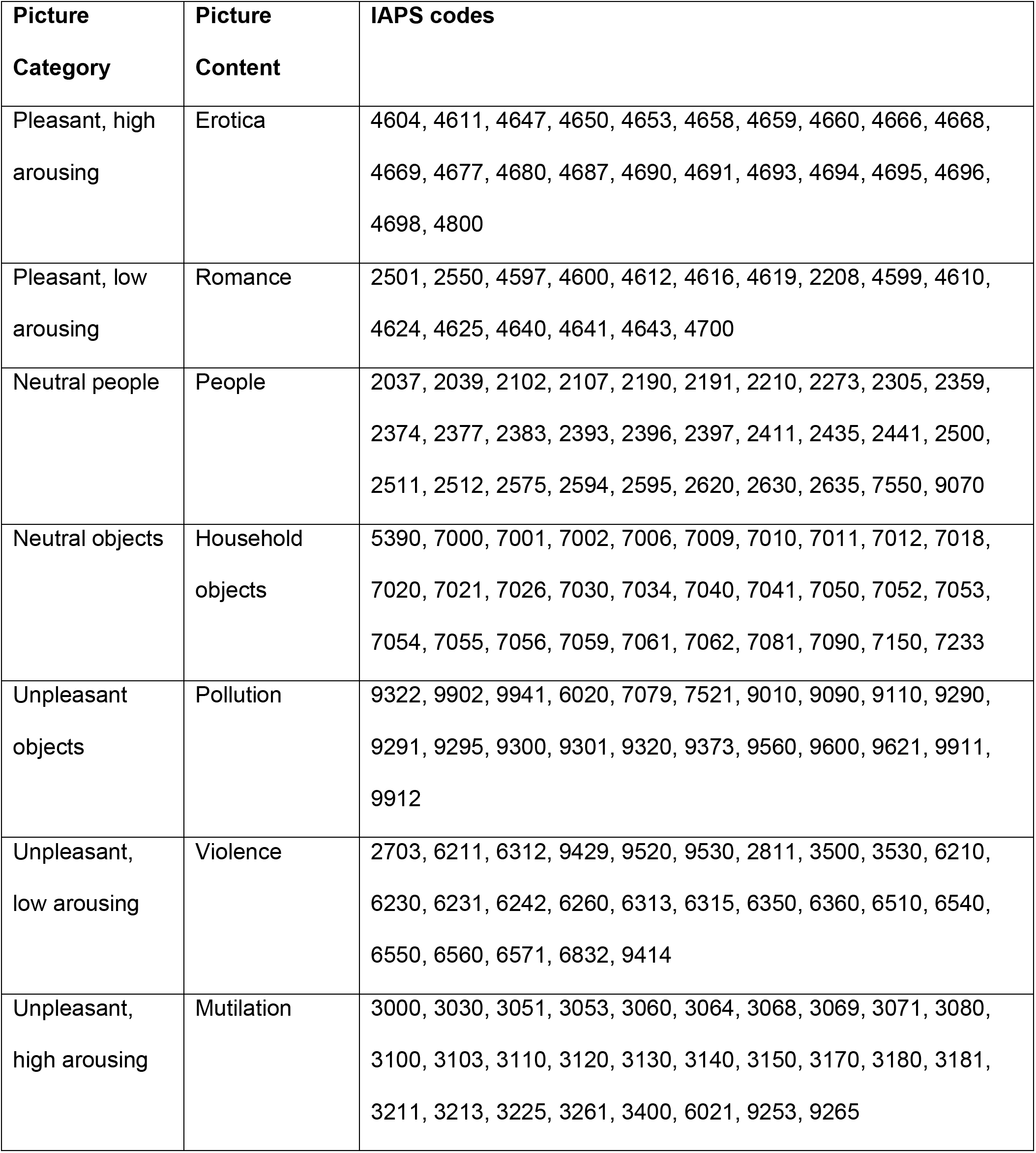
A list of the IAPS images used in the experiment by picture category and content.

**Supplementary Fig. S1.**
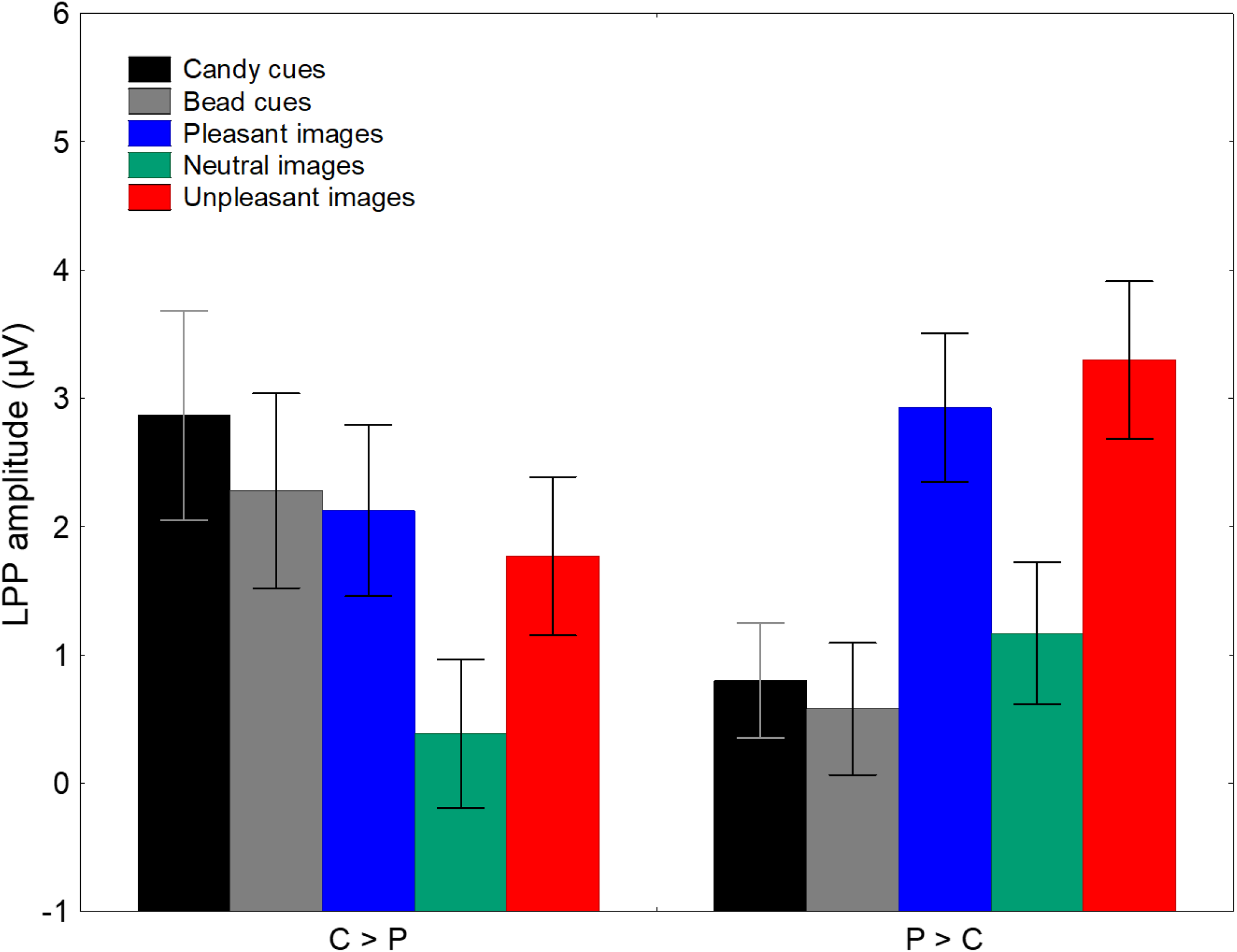
Average LPP responses for each parent picture category (Pleasant, Unpleasant, etc) by LPP group. After averaging LPP amplitudes for the emotional image subcategories (pleasant: erotica and romance; unpleasant: mutilations, violence, and pollution) together, we found that the C>P group had higher LPP amplitudes for food cues than for pleasant images, whereas the P>C group had higher LPP amplitudes for pleasant images than for food cues. Error bars: 95% CIs.

**Supplementary Fig. S2.**
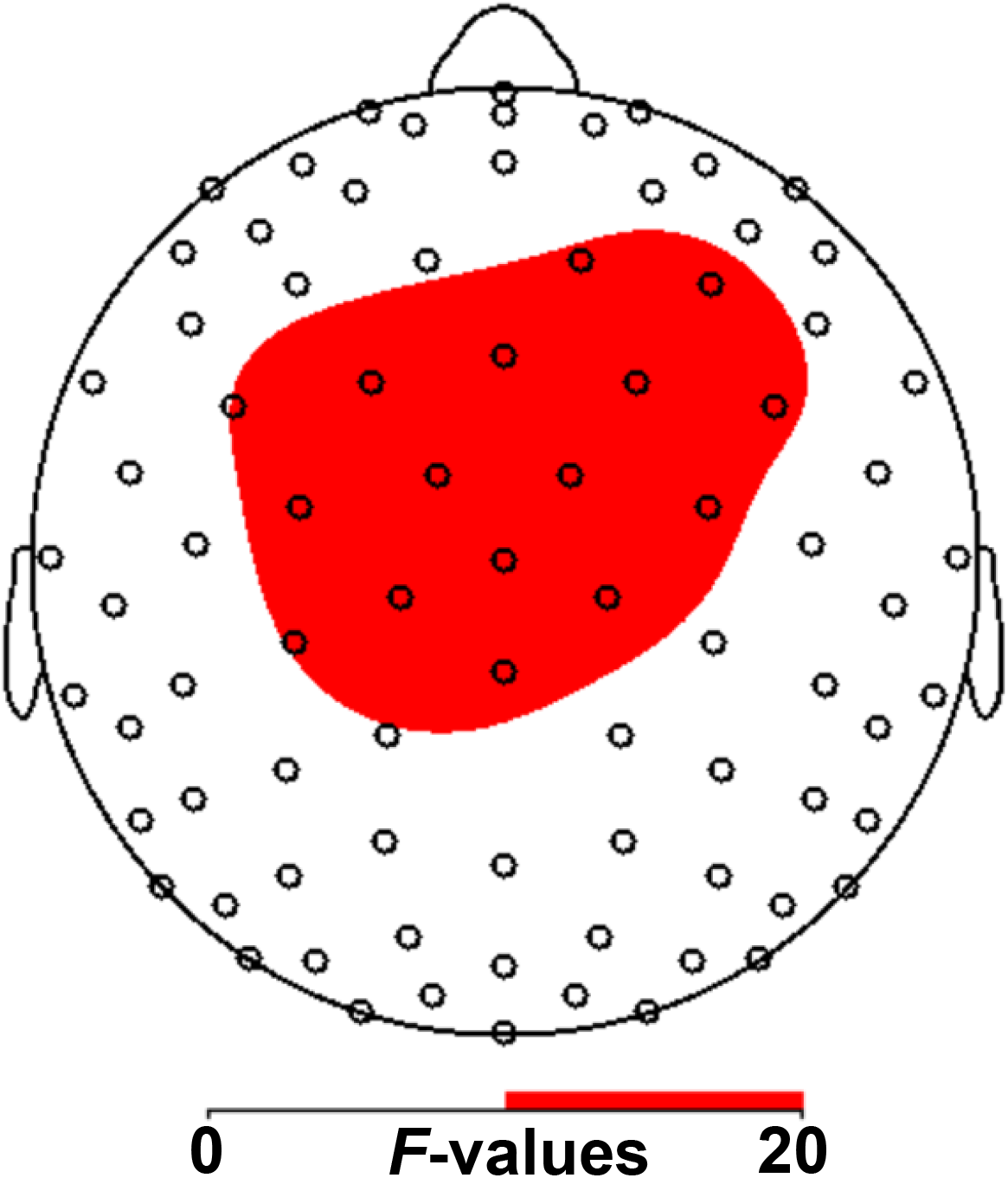
After conducting a repeated-measures ANOVA comparing the theta power under the candy, bead, and neutral conditions for each time point and EEG sensor, we identified the *F*-values that were statistically significant during the 0- to 200-ms time window, during which theta effects are usually the highest. Shown are the EEG sensors that were statistically significant during the 0- to 200-ms time bin based on a Bonferroni correction. The color bar in red indicates the *F*-value magnitude. Critical *F* ≈ 10.

**Supplementary Fig. S3.**
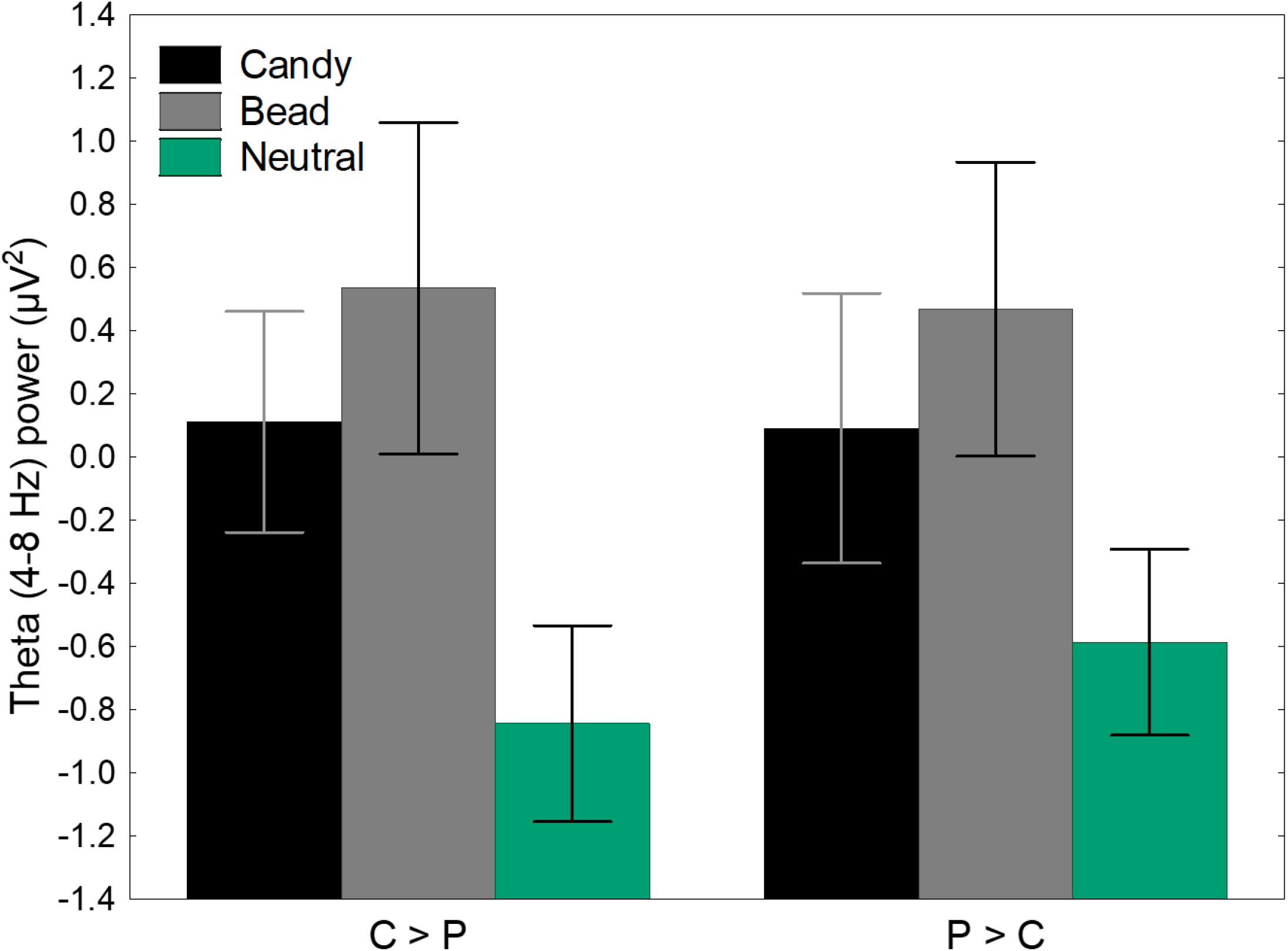
Average theta power from the pooled midfrontal sensors in the 0-200 ms time bin under the candy, bead, and neutral conditions by C>P and P>C groups. We compared the theta power for the candy, bead, and neutral conditions in the participant groups formed using *k*-means clustering with LPP data. We did not find significant differences in theta power in the C>P and P>C groups.

